# Bayesian analysis of isothermal titration calorimetry for binding thermodynamics

**DOI:** 10.1101/327676

**Authors:** Trung Hai Nguyen, Ariën S. Rustenburg, Stefan G. Krimmer, Hexi Zhang, John D. Clark, Paul A. Novick, Kim Branson, Vijay S. Pande, John D. Chodera, David D. L. Minh

**Author notes:** **For correspondence:** (JDC); (DDLM). These authors contributed equally to this work (Trung Hai Nguyen, Ariën S. Rustenburg). ^‡^Nurix, San Francisco, CA 94158, USA. ^§^Genentech, South San Francisco, CA 94080, USA.

## Abstract

Isothermal titration calorimetry (ITC) is the only technique able to determine both the enthalpy and entropy of noncovalent association in a single experiment. The standard data analysis method based on nonlinear regression, however, provides unrealistically small uncertainty estimates due to its neglect of dominant sources of error. Here, we present a Bayesian framework for sampling from the posterior distribution of all thermodynamic parameters and other quantities of interest from one or more ITC experiments, allowing uncertainties and correlations to be quantitatively assessed. For a series of ITC measurements on metal:chelator and protein:ligand systems, the Bayesian approach yields uncertainties which represent the variability from experiment to experiment more accurately than the standard data analysis. In some datasets, the median enthalpy of binding is shifted by as much as 1.5 kcal/mol. A Python implementation suitable for analysis of data generated by MicroCal instruments (and adaptable to other calorimeters) is freely available online.

## Introduction

Isothermal titration calorimetry (ITC) is a widely used biophysical technique for measuring the binding affinity between small molecules and biological macromolecules (such as proteins and RNA [8, 15, 26, 27]), as well as between proteins [37]. In addition to simple two-component (one-to-one) binding processes, ITC may also be used to study more complex processes such as competitive binding [15, 36], binding cooperativity [2], and binding events coupled to changes in the protonation state [6, 28] or tautomeric state [11] of one or more components. Provided reaction rates are slower than cell mixing times, ITC can even be used to study the kinetics of association [23].

Here, we focus on the thermodynamics of simple two-component association (one-to-one binding). A unique and powerful property of ITC is that it can not only determine the free energy of binding (Δ*G*), but also decompose it into enthalpy (Δ*H*) and entropy (Δ*S*) without having to resort to multiple experiments at different temperatures to determine these quantities via the van’t Hoff equation. This decomposition has been used to draw conclusions into, for example, how entropy is related to antibody flexibility [35] and ordering of disordered loops [4] during antibody affinity maturation. It has also been used to suggest that iterative improvements in generations of drugs result in their interactions being increasingly driven by enthalpy [14]. Furthermore, it has been used to suggest how force fields might be improved [9].

It is possible to perform enthalpy-entropy decomposition with ITC because the instrument not only detects a binding process, but can determine the heat of binding. The raw data from an ITC instrument is the differential power required to maintain a reference cell at the same temperature as the *titrand* in a sample cell (usually a macromolecule dissolved in buffer) as a *titrant* (usually a small molecule ligand) is injected into it. The experimental data 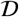 can be summarized as the measured heats of injection, 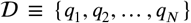 obtained by integrating the differential power over the duration of each injection. Thermodynamic parameters are then determined by fitting binding heat models (expressions for the heat in terms of unknown thermodynamic and experimental parameters) to the integrated heat [38]. The standard protocol for parameter estimation, implemented in the Origin software package [17] distributed with the popular MicroCal VP-ITC instrument [18], uses a nonlinear least squares fit to estimate the association constant *K*_*a*_, enthalpy Δ*H*, and stoichiometry *n* (number of binding sites per mole of receptor), along with their estimated uncertainties.

Unfortunately, this established procedure for analyzing ITC data does not accurately determine uncertainties for enthalpy-entropy decomposition because it fails to account for all relevant sources of error. In a large-scale interlaboratory study (ABRF-MIRG’02) of a model protein: small molecule binding reaction — the binding of carboxybenzenesulfonamide (CBS) to bovine carbonic anhydrase II (CAII) — the variation among the reported ITC binding constant and enthalpy from 14 participants was more than an order of magnitude larger (and up to *three* orders of magnitude larger) than standard errors reported by the individual least squares analyses [21].

Spectrophotometric results suggested that titrant concentration errors were likely a major cause of this unexpectedly large variation. The standard analysis method accounts for error in the titrand concentration by treating the stoichiometry *n* as a free parameter that can take any real and positive value. On the other hand, the titrant concentration, likely an important source of discrepancies among laboratories [34], is often treated as exactly known. While precise titrant concentrations are systematically achievable [1], strong evidence suggests that large (10-20%) errors in titrant concentration are widespread even amongst laboratories skilled in biomolecular calorimetry [21]. It is possible to explicitly treat titrant concentration error in nonlinear least squares fitting [1], but this is not typically performed.

In addition to concentration error, another important source of error that is frequently neglected is the so-called *first injection anomaly*, in which the heat of injection from the first injection is smaller than expected. The anomaly often emerges due to backlash in the motorized screw mechanism used to drive the syringe plunger [20]; if the last operation of the plunger prior to the first injection is upwards, then less titrant will be injected via a subsequent downward movement of the plunger. This issue may be overcome by executing a short downward movement of the plunger prior to insertion into the sample cell. Another contributing factor to the first injection anomaly is leakage of titrant out of the syringe during instrument equilibration. Because the initial injection generally carries the largest magnitude of heat per mole of titrant injected, the first injection anomaly (or the inability to account for it) can lead to significant errors in reported measurements.

Here we introduce a new data analysis protocol that accounts for these sources of error and, as we shall show, more accurately estimates the uncertainty in derived thermodynamic parameters — especially entropy and enthalpy. The approach is modular; additional sources of uncertainty or variability can be modeled through simple extensions of the model. Importantly, this analysis procedure also allows the joint uncertainties in entropy and enthalpy to be resolved, an essential requirement to evaluating hypotheses regarding entropy-enthalpy compensation. Our approach is based on Bayesian statistics, which uses the *posterior* probability distribution,

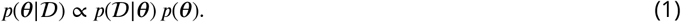

where 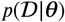 is the *likelihood*, a conditional probability of observing data 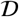 (in our case, the injection heats {*q*_1_,…,*q*_*N*_}) given unknown thermodynamic parameters *θ*. *p*(*θ*) is the *prior* probability, a function describing foreknowledge of the parameters *θ* before conditioning this distribution on the observed data 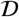 from this experiment.

A Bayesian analysis has several significant potential advantages over the standard analysis protocol, including:

1. **Multimodal posteriors**: Bayesian analysis makes no assumptions about the shape of the posterior. Therefore, it can treat multimodal posteriors in which two or more distinct sets of parameters describe the data. On the other hand, the standard analysis assumes a multivariate Gaussian, which is based on a single mode.
2. **Nonlinear parameter correlation**: It is feasible to determine whether parameters are correlated, even if correlations are nonlinear.
3. **Modularity**: Additional sources of uncertainty can be incorporated in a modular fashion simply by adding more random variables (nuisance parameters) with associated priors.
4. **Integration of multiple experiments**: It is possible to incorporate information from multiple measurements and even from multiple experimental techniques. The posterior probability of a parameter is simply the product of posteriors for each measurement. Information from control experiments, such as a blank titration or prior standard measurements, can be incorporated into the prior.
5. **Optimal experimental design**: New experiments that maximize the gain of new information can be automatically identified. By using techniques from Bayesian experimental design [3], one can choose among many potential experiments those that would maximize the gain of *new* information, either sequentially or in batches.

To clarify, it is possible for analyses based on nonlinear regression to integrate some of these features. In nonlinear regression, parameter distributions are inherently non-Gaussian and two-dimensional contour plots of different parameters may have non-ellipsoid shapes, indicating nonlinear correlations [29, 33]. It is also possible to integrate multiple experiments with a global fit. However, these features are not available in the standard protocol.

Recently, Duvvuri et al. [7] described a new python package for the Bayesian analysis of ITC experiments. For the analysis of single experiments, their results were consistent with Origin. They were also able to integrate data from multiple buffers, titrant/titrand ratios, and temperatures. However, they did not perform substantial error analysis.

Our present work is based on a different new python package and we more carefully consider the uncertainty of different analysis protocols. The primary criterion we use to evaluate and compare analysis protocols is based on interval estimates. Interval estimates have somewhat different meanings in Bayesian and frequentist statistics. In frequentist statistics, the *α*% confidence interval is expected to contain the true value *α*% of the time. A confidence interval is inaccurate if the percentage of estimated intervals that contain the true value deviates from *α*%. In Bayesian statistics, the *credible* interval is not necessarily intended to contain the true value a specific percentage of the time; it is simply a region that contains *α*% of the posterior probability. Nonetheless, for the purposes of comparing uncertainties, we evaluate whether the Bayesian credible interval (BCI) obtained from our model serves as an accurate confidence interval compared to the confidence interval from the standard nonlinear regression protocol (NlRCI). Previously, BCIs have been shown to work well as confidence intervals for binding thermodynamics and reference scattering patterns in analyses of X-ray scattering experiments of protein:ligand binding [19].

## Experimental

### Titration of Mg(II) into EDTA

In order to assess the effectiveness of the Bayesian approach in describing the true uncertainty in the experimental measurements, we studied a simple complexation reaction—the 1:1 binding of Mg(II) to the chelator EDTA—for which multiple experimental replicates can be easily collected. The entire ITC experiment was repeated *from scratch*—with all solutions prepared completely independently so that any concentration errors would be fully independent—a total of 14 times. This is critical, as simply repeating the experimental measurement with the same stock solutions would not capture the true experimental variability. For each trial, the titrant (MgCl_2_), titrand (EDTA), and buffer (50 mM Tris-HCl pH 8.0) were weighed and dissolved to prepare solutions at the two planned concentrations for the titrant MgCl_2_ and the titrand EDTA. In the first five trials, we prepared the titrant and titrand concentrations as 1.0 mM and 0.1 mM, respectively. In the other nine trials, the titrant and titrand concentrations were prepared as 0.5 mM and 0.05 mM, respectively.

Magnesium chloride hexahydrate [MgCl_2_·(H_2_O)_6_] was purchased from Fisher Scientific (Catalog No. BP214-500, Lot No. 006533) and anhydrous ethylenediaminetetraacetic acid (EDTA) was purchased from Sigma-Aldrich (Catalog No. E6758-500G, Batch No. 034K0034). Tris base was purchased from Fisher Scientific (Catalog No. BP154-1, Lot No. 082483). Buffer was prepared by weighing Tris base, adding MilliQ water, and adjusting the final pH to 8.0 by dropwise titration with HCl or NaOH. Solutions were prepared by weighing powder and adding the appropriate amount of buffer, neglecting the volume occupied by powder, to make a concentrated solution (15 mM for MgCl_2_ and 1.0 mM for EDTA). To maximize the number of significant figures, at least 0.1 g of MgCl_2_ and 0.01 g of EDTA were weighted out. The solutions were then further diluted with buffer to prepare the titrant and titrand. For example, to prepare a 0.1 mM solution of EDTA, a pipetman was used to measure 9 parts buffer to 1 part of 1.0 mM EDTA.

ITC measurements were performed on a MicroCal VP-ITC calorimeter. The experiments consisted of a total of 24 injections, with the first injection programmed to deliver 2 *μ*L of titrant (MgCl_2_) into the sample cell, and the remaining 23 injections programmed to deliver 12 *μ*L. Data was collected for 60 s prior to the first injection and 300 s for each injection. The injection rate for all injections was 0.5 *μ*L/s. All experiments were conducted at 298.1 K, and the reference power was fixed at 5 *μ*cal/s.

The baseline was corrected and injection heats integrated using NITPIC [12].

### Titration of phosphonamidate-type inhibitors into thermolysin

To demonstrate our approach on protein:ligand systems, we also analyzed titrations of phosphonamidate-type inhibitors into thermolysin initially described in Krimmer et al. [13]. For each individual measurement, lyophilized thermolysin powder was freshly weighed (1.5-2 mg) and dissolved in an appropriate volume of buffer to achieve a concentration of 30 *μ*M. The concentration was confirmed by ultraviolet absorption at 280 nm. Prior to measurement, the thermolysin soluton was centrifuged for 8 min at 8150 g. In contrast, one solution was prepared for all measurements with each ligand by dissolving the pure powder (0.3-0.4 mg) in buffer without the addition of DMSO. A MX5 microbalance from Mettler Toledo (Switzerland) with a readability of 1 *μ*g and a repeatability of 0.8 *μ*g was used for the sample weighting. Measurements were repeated in this fashion at least nine times. Results reported in Krimmer et al. [13] were based on three repetitions with a fresh batch of thermolysin and after optimizing ITC parameters. In contrast, our present analysis was based on all available data for each system except for a small subset with a large baseline shift in the middle of an injection.

Lyophilized thermolysin (EC number 3.4.24.2) from *Bacillus thermoproteolyticus* was purchased from Calbiochem (EMD Biosciences). The inhibitors (Figure 1) P-((((benzyloxy)carbonyl)amino)methyl)-N-((S)-4-met hyl-1-oxo-1-(propylamino)pentan-2-yl)phosphonamidicacid (ligand **1**), P-((((benzyloxy)carbonyl)amino)methyl)-N-((S)-1-(isobutylamino)-4-methyl-1-oxopentan-2-yl)phosphonamidicacid (ligand **2**), and P-((((benzyloxy)car bonyl)amino)methyl)-N-((S)-4-methyl-1-(((S)-2-methylbutyl)amino)-1-oxopentan-2-yl)phosphonamidicacid (ligand **3**), were synthesized as previously described [22]. (Crystal structures of ligands **1** (PDB ID 4MXJ), **2** (PDB ID 4MTW), and **3** (PDB ID 4MZN) in complex with thermolysin have been previously reported [13]). All measurements were performed with a buffer composed of 20 mM HEPES (pH 7.5), 200 mM NaSCN, and 2 mM CaCl_2_.6H_2_O. HEPES was purchased from Carl Roth (Catalog No. 9105.3, Batch no. 192184596), NaSCN was purchased from Fluka Analytical (Catalog No. 71938-1KG, Lot no. BCBC9384V), and CaCl_2_·6H_2_O was purchased from Carl Roth (Catalog No. T886.2, Lot no. 433205269). Prior to measurement, the buffer was filtered through a 0.22 μm filter and degassed under reduced pressure.

ITC measurements with thermolysin were performed on an MicroCal ITC_200_ calorimeter from GE Healthcare (Piscataway, New Jersey). After an initial delay 170 or 180 sec, the initial injection (0.3-0.5 *μ*L) was followed by 19-26 main injections (1.2-1.5 *μ*L). The duration of the injection (in sec) was twice the value of the volume (in *μ*L). All measurements were performed at a temperature of the measurement cell of 298.15 K, a stirring speed of 1000 rpm, titrand (thermolysin) concentration of 30 *μ*M, and a titrant (ligands **1-3**) concentration of 400 *μ*M. For details on each protocol, see Tables S1-S3 of the Supplementary Material.

As with the Mg(II):EDTA data, the baseline was corrected and injection heats integrated using NITPIC [12].

**Figure 1.**
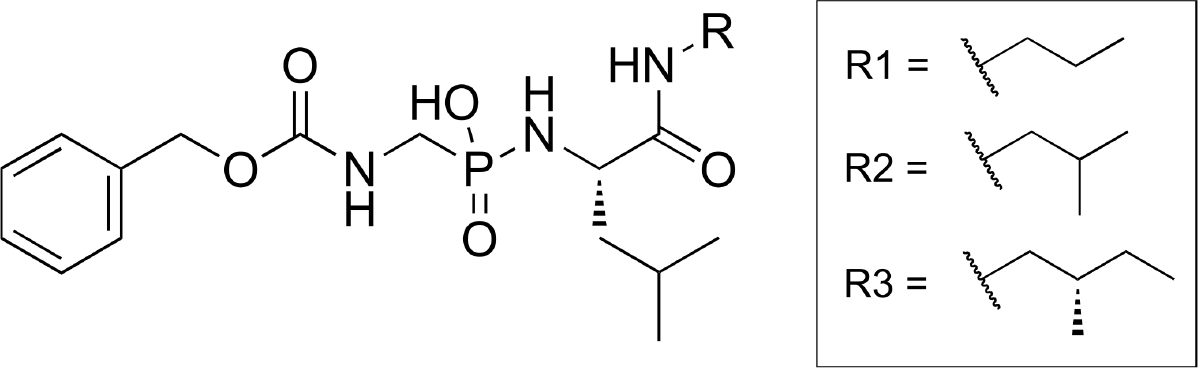
Chemical structures of thermolysin ligands used in this study.

### CBS:CAII dataset from the ABRF-MIRG’02 study

Finally, we considered a protein:ligand ITC dataset from a previously published study which demonstrated large interlaboratory variation far in excess of reported error estimates [21]. Injection heat data were digitized from Figure 4 in the ABRF-MIRG’02 paper [21], which includes 14 ITC datasets measured fully independently on identical source material (aliquots of CAII and dry powder stocks of CBS) by independent laboratories. Dataset 2 was generated by an instrument called the CSC 4200 ITC (see Table 2 in [21]) for which we could not find the user’s manual to obtain information such as the cell volume. Therefore, we excluded this dataset. We also excluded dataset 4 because we were not able to reliably digitize the large number of injections. For other datasets, the experimental design parameters were taken from Table 2 of the study, while the reported thermodynamic parameters and standard errors were taken from Table 3 [21]. In the ABRF-MIRG’02 study [21], most experiments obtained standard errors were using a nonlinear least squares fit. The exceptions were datasets 10 and 14, in which the standard deviation was obtained by repeating the same experiment 3 and 5 times, respectively. In these datasets, it was not clearly specified whether the entire experiment or just the titration was repeated in each replicate.

### Frequentist confidence intervals

Origin software was used to perform nonlinear least squares fit of the heat data to obtain the binding constant *K*_*a*_, enthalpy Δ*H*, and the stoichiometry number *n*, and their corresponding standard errors. Each parameter was assumed to be normally distributed and the standard error was used as a standard deviation. The lower and upper bounds of the *α*% confidence interval were the 1 − *α*/2 and 1 + *α*/2 percentile, respectively, of the normal distribution with a mean as the point estimate and standard deviation as the reported uncertainty.

### Sampling from the Bayesian posterior

Our Bayesian model is constructed to infer the unknown true parameters,

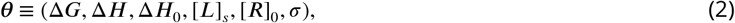

which represent

- Δ*G*: the free energy of binding
- Δ*H*: the enthalpy of binding
- Δ*H*_0_: the enthalpy of dilution and stirring per injection
- [*L*]_*s*_ and [*R*]_0_: the concentrations of titrant in the syringe and of the titrand in the cell, respectively
- *σ*: the standard error of heat measurement per injection, a nuisance parameter needed to write the data likelihood. The model assumes all injections include the same number of power measurements.

#### Likelihood

The data 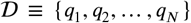 consists of the observed heats per injection determined by integrating the differential power over the injection time. The corresponding data likelihood function was based on the assumption that, because the *observed* injection heat *q*_*n*_ is the sum of many power measurements, the measurement error added to the *true* (unknown) heat 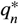 will be normally distributed due to the central limit theorem,

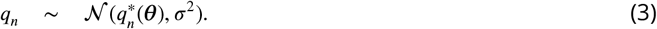

The total data likelihood for 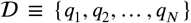 is therefore given by

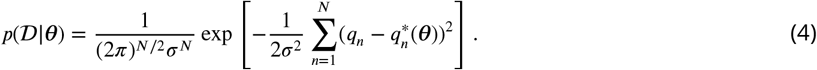

The model heats 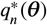 are a function of the parameters *θ*. See Appendix for details of the binding model relating *θ* to the true heats 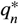.

#### Priors

The prior *p*(*θ*) was a product of priors for each parameter, *p*(*θ*) = Π*_j_ p*(*θ_j_*). Uniform priors were chosen for Δ*G*, Δ*H*, and Δ*H*_0_:

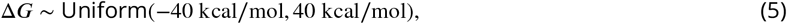

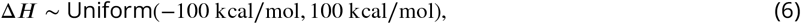

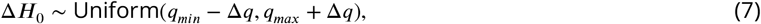

where *q*_*min*_ = min{*q*_1_,*q*_2_, … *q*_*N*_}, *q*_*max*_ = max{*q*_1_,*q*_2_, … *q*_*N*_} and θ*q* = *q*_*max*_ − *q*_*min*_, usually reported in units of cal.

We used three different sets of priors for the true concentrations of titrant in the syringe, [*L*]_*s*_, and receptor in the cell, [*R*]_0_ (Table 1): General, Flat [*R*]_0_, and Comparison. All of the concentration models make use of the fact that concentrations must be positive. In the General model, both concentrations are assigned lognormal priors with the mean and standard deviation given by their stated experimental values and corresponding experimental uncertainties due to preparation steps,

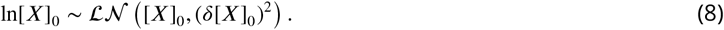

In the absence of specific quantification of the titrant concentration uncertainty, we assumed a value of *δ*[*X*]_0_ equal to 10% of the provided [*X*]_0_. This specified uncertainty is in line with quantification of typical laboratory titrant concentration errors observed by Myszka et al. [21]. In cases where the practitioner uses an orthogonal method to quantify titrant concentration or carefully tracks the uncertainty during preparation steps, as described in Boyce et al. [1], this more precise concentration uncertainty could be used instead. Alternatively, *δ*[*X*]_0_ could be treated as a free nuisance parameter. Although the parameter may not be precisely determined from a single ITC experiment, it could potentially be elucidated by sampling from a Bayesian posterior based on multiple measurements with the same titrand solutions, analogous to a global fitting procedure possible with nonlinear least squares [32].

In the Flat [*R*]_0_ model, a uniform prior was used for [*R*]_0_ such that,

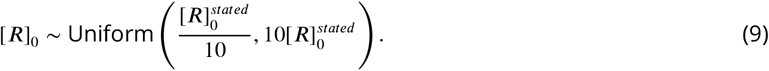

This model is useful in cases where the receptor concentration is not clearly known, such as when the sample is impure or partially degraded. Due to potential degradation of protein used in some ITC measurements, we used this model in our analysis of data for thermolysin. Finally, in the Comparison model, we used a uniform prior for [*R*]_0_ and a sharply peaked prior for [*L*]_*s*_ such that 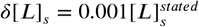.

**Table 1.**
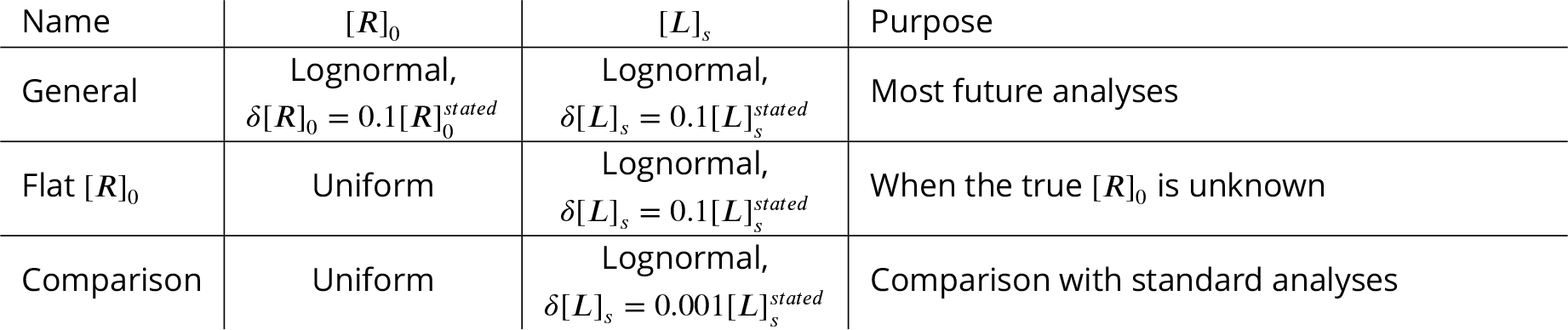
Summary of concentration priors used in this manuscript.

The Comparison model mimics the treatment of concentrations in standard nonlinear least squares fitting. In the standard procedure, [*L*]_*s*_ is assumed to be precisely the stated value while [*R*]_0_ can take any positive value that minimizes the total residual sum of squares. There is no penalty for changing [*R*]_0_ from its stated value. This is consistent with flat prior for [*R*]_0_ a sharp prior for [*L*]_*s*_. On the other hand, the General model allows for but penalizes deviations from the stated values. In the absence of further information, we believe that the General model is the most justified of the three models because concentrations are likely to be close to their stated values. Our main reason for performing calculations with the Comparison model was to isolate the effects of concentration models from other aspects of the Bayesian analysis.

Finally, since even its order of magnitude may be unknown, an uninformative Jeffreys prior [10] was assigned to the noise parameter *σ*,

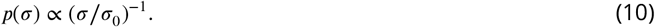

where *σ*_0_ ≡ 1 cal is a reference quantity that simply renders the ratio *σ*/*σ*_0_ dimensionless. This model assumes that the injection heat measurement uncertainty *σ* is constant for all injections. This may be a good approximation when the same number of power measurements are integrated for each injection (i.e., when injections are of identical duration), but when experiments contain injections of different durations, the noise variance *σ*^2^ should be proportional to the number of power measurements summed to give the injection heat (with all other things being held constant). More complex noise variance models (such as those considered in [31]) could also be considered. The noise model could also be improved using calibration experiments based on the same protocol (such as blank titrations), or even other data collected on the instrument for other systems; in these cases, likelihoods from independent experiments are simply multiplied.

While we used uninformative priors (except for our concentrations) in this study, alternative priors for other parameters can be used. If some knowledge of thermodynamic parameters or concentrations is available from another type of experiment, e.g., spectrophotometric measurements, then these can be incorporated into their respective priors. In such cases, the prior could be normally-distributed with the sample mean and standard deviation as parameters. Another way to parameterize the prior for concentrations is by careful propagation of error during the sample preparation process (from estimates of known pipetting error magnitudes, known analytical balance accuracies, and reported compound purities). Alternatively, the posterior from a previous (e.g., pilot) ITC experiment can be used as the prior to integrate the information from a second ITC experiment with different experimental parameters.

#### Sampling from the posterior

Because it is complex and multidimensional, the posterior distribution (Eq. 1) is not amenable to direct sampling using acceptance-rejection or another method that generates independent and identically distributed variates. To compute statistics such as the mean, median, mode, credible intervals, and marginal distributions of the posterior, we instead sampled from the posterior using Markov chain Monte Carlo (MCMC) [16] simulation. Initial values were chosen as follows:

- for [*L*]_*s*_ and [*R*]_0_, the stated (intended) concentration was used
- for Δ*H*, Δ*G*, and Δ*H*_0_, initial values of zero (in their appropriate energy units) were used
- for *σ*, the standard deviation of the last four injection heats was used as an initial guess

Parameters were updated by sequential Gibbs sampling where each parameter was updated via Metropolis-Hastings sampling:

1. For each single parameter, a proposal is drawn from a normal distribution centered at the current value, and a scale of unity for Δ*H*, Δ*G*, and Δ*H*_0_, or the initial guess value for *σ*, [*L*]_*s*_, and [*R*]_0_;
2. The trial move is accepted or rejected according to the Metropolis criterion. If it is accepted, the next value in the Markov chain is the trial move. If it is rejected, the next value in the Markov chain is the original value.

MCMC was performed using a python library that we wrote, bayesian-itc (https://github.com/choderalab/bayesian-itc). bayesian-itc uses the Metropolis-Hastings implementation in the PyMC [24] library to perform MCMC sampling. For each experiment, sampled parameters were stored after every 2000 MCMC trial moves for a total of 5000 samples. Each sample from the Bayesian posterior is a set of six values, as described in Equation 2. For each parameter, the *α*% BCI is estimated based on the shortest interval that contains *α*% of the MCMC samples.

The precise version of library used in this manuscript was committed to github on May 2, 2018 at https://github.com/nguyentrunghai/bayesian-itc/tree/d8cbf43240862e85d72d7d0c327ae2c6f750e600. The directory entitled `analysis_of_Mg2EDTA_ABRF-MlRG02_Thermolysin` contains all the data needed to reproduce the figures in this manuscript.

### The Kullback-Leibler divergence quantifies differences between thermodynamic parameter distributions obtained from Bayesian and nonlinear least-squares approaches

To compare posterior marginal distributions, we computed the Kullback-Leibler divergence (KL-divergence), between the posterior marginal densities in the two most important thermodynamic quantities of interest, (Δ*G*, Δ*H*),

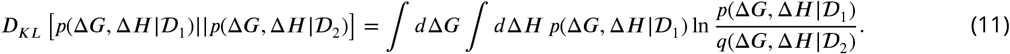

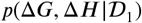 and 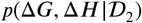 are the posterior marginal densities specified by two different experiments with associated datasets 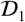 and 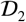. This metric, commonly used as a measure of deviation between two probability densities, can be interpreted as the amount of information lost when 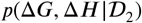 is used to approximate 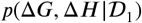. The marginal posterior density for each experiment was estimated by using a Gaussian kernel density estimate (KDE) based on MCMC samples for Δ*G* and Δ*H* (ignoring other parameters). We used the `KernelDensity` package implemented in scikit-learn [25] to estimate the density *p*(Δ*G*, Δ*H*). The bandwidth for Gaussian kernel was set to 0.03 kcal/mol. Although the Kullback-Leibler divergence can be analytically computed for Gaussian densities, we also used the same KDE method to estimate probability densities *p*(Δ*G*, Δ*H*) for nonlinear regression. Samples for Δ*G* and Δ*H* were drawn from Gaussian distributions with the mean and standard deviation based on nonlinear regression point estimates and errors, respectively.

## Results and Discussion

### MCMC sampling leads to precise estimates of Bayesian credible intervals

Our MCMC sampling protocol appears to yield precise estimates of 95% BCIs (Figure 2 and Figures S1 to S4 in the Supplementary Material). In all of the selected systems, the estimated 95% BCIs do not substantially change after considering about 2000 samples. The standard deviation of estimated upper and lower bounds over the five independent simulations in each system was less than 5% of the length of the average interval. Therefore, we are confident that the number of MCMC samples and mixing of the MCMC chain is sufficient to yield consistent estimates of the BCIs and other statistics of interest.

### Bayesian ITC yields unimodal distributions of linearly correlated parameters

Bayesian analysis permits multimodal posteriors and nonlinear parameter correlations to be investigated. Qualitative trends in the posterior density may be visualized by generating histograms of MCMC samples drawn from the posterior. For our systems, representative 1 D marginal distributions of key parameters (Figure 3) are unimodal. Although some skew is evident in Δ*H*, the Gaussian distribution could be considered a reasonable approximation for most of these parameters. Our observation is consistent with previous analyses of nonlinear regression which showed that a Gaussian assumption is appropriate when the magnitude of statistical error is less than 10% of a parameter [29, 33].

**Figure 2.**
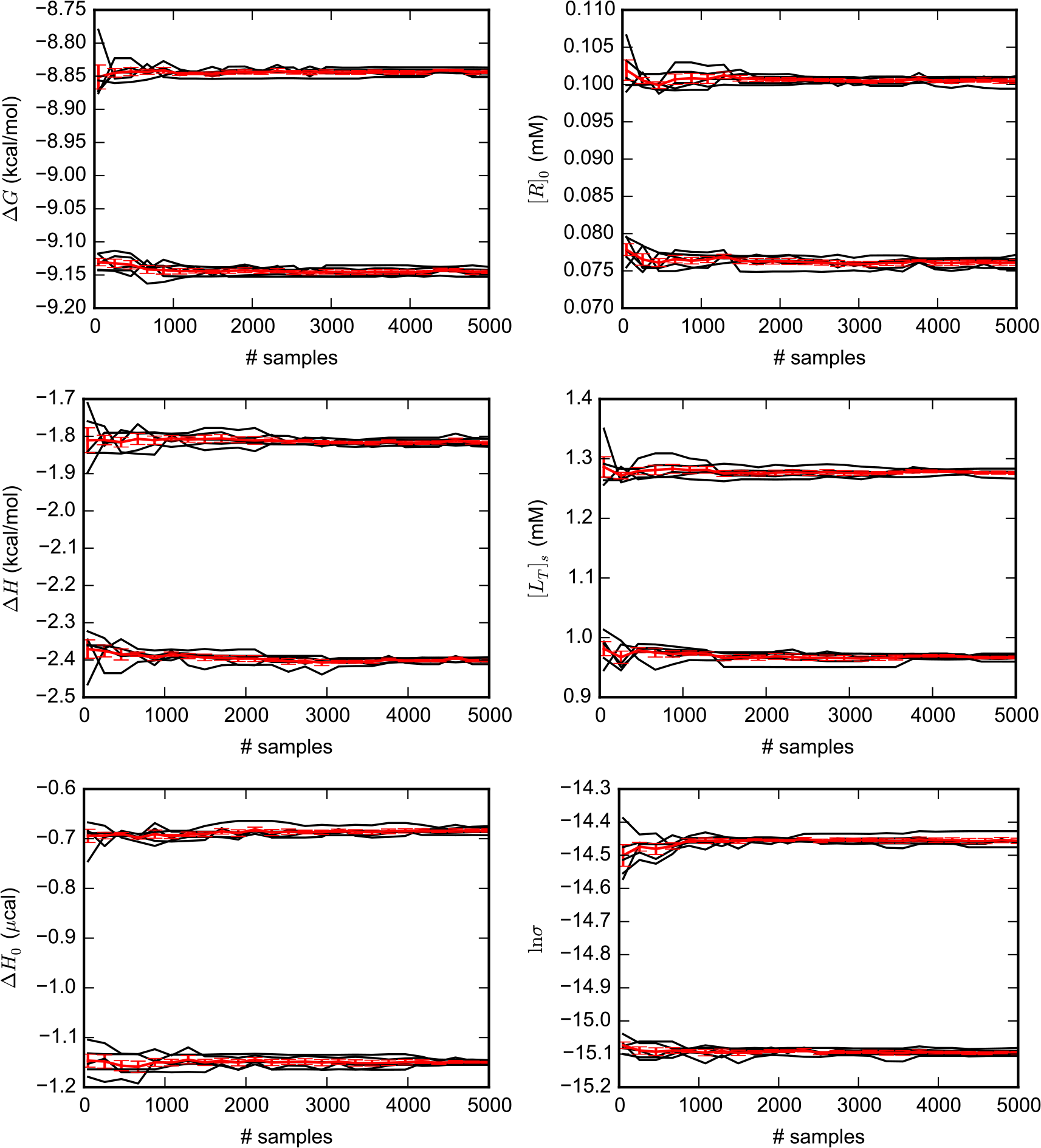
Convergence of 95% Bayesian credible intervals (BCIs) with MCMC sampling. 5000 MCMC samples were generated from the Bayesian posterior (General model) for several variables based on one ITC experiment measuring Mg(II):EDTA binding. For five independent repetitions of the MC simulations, the black lines are running estimates, as the number of samples is increased, of the upper and lower limits of 95% BCIs. The red line and error bars are the average and standard deviation across the five independent simulations. Similar plots for ligands **1-3** binding to thermolysin and CBS:CAII are available as Figures S1 to S4 in the Supplementary Material.

Representative 2D marginal distributions (Figure 4) show that some pairs of parameters are nearly independent and others are highly correlated, with varying degrees of correlation in between. Of particular interest is the fact that while the free energy Δ*G* and enthalpy Δ*H* are mostly uncorrelated (top left of Figure 4), there is high correlation between the enthalpic (Δ*H*) and entropic (*T*Δ*S*) contributions to binding (top right of Figure 4) and between Δ*H* and the receptor concentration [*R*]_0_ (bottom right of Figure 4). These correlations are not considered in the standard nonlinear regression analysis.

Given that the correlations appear to be linear, they can be succinctly summarized via the correlation coefficient. The estimated correlation matrix shown in Table 2 indicates that the titrant [*L*]_*s*_ and titrand concentrations [*R*]_0_ are highly correlated with each other and with the enthalpy Δ*H* but only weakly with Δ*G*. This result is consistent with Tellinghuisen [30], who evaluated the sensitivity of the binding constant and enthalpy to changes in concentration.

**Table 2.** **Correlation matrix** estimated from the Bayesian posterior (General model) for an Mg(II):EDTA binding dataset. Numbers in parentheses denote the uncertainty in the last digit.

| | $\Delta G$ | $\Delta H$ | $\Delta H_0$ | $[L]_s$ | $[R]_0$ | $\ln \sigma$ |
| --- | --- | --- | --- | --- | --- | --- |
| $\Delta G$ | 1 | 0.50(1) | 0.280(8) | 0.541(9) | 0.553(9) | 0.00(2) |
| $\Delta H$ | 0.50(1) | 1 | -0.07(1) | 0.9893(2) | 0.9884(2) | 0.01(1) |
| $\Delta H_0$ | 0.280(8) | -0.07(1) | 1 | 0.00(1) | 0.01(1) | 0.016(9) |
| $[L]$ | 0.551(9) | 0.9893(2) | 0.00(1) | 1 | 0.998 87(3) | 0.01(1) |
| $[R]$ | 0.553(9) | 0.9884(2) | 0.01(1) | 0.998 87(3) | 1 | 0.01(1) |
| $\ln \sigma$ | 0.00(2) | 0.01(1) | 0.016(9) | 0.01(1) | 0.01(1) | 1 |

Estimates of concentrations and Δ*H* are correlated because the effect of changing one of the parameters can be largely counteracted by changing another. When samples from a Bayesian posterior for MG(II):EDTA binding were used to parameterize a simple linear model for [*L*]_*s*_ and Δ*H* as a function of [*R*]_0_, different parameter values led to essentially the same integrated heat curve (Figure 5). An important implication of this enthalpy-concentration compensation is that given a measured integrated heat curve, the precise values of the three parameters are underdetermined; by itself, ITC cannot simultaneously determine the titrant or titrand concentration and the enthalpy of binding.

### Median enthalpy estimates are sensitive to the titrand concentration model

Even though nonlinear least squares fitting and Bayesian analysis are based on the same binding model, other variations in the analysis procedure may lead to different estimates of Δ*G* and Δ*H*. We compare different analysis methods by considering how the median (which is less sensitive to outliers than the mean) of each quantity within a dataset. For all datasets, the median estimate of Δ*G* is largely consistent across the different analysis methods. In contrast, with the thermolysin datasets, Δ*H* estimates are consistent between all models except for the General model, which differ by as much as 1.5 kcal/mol (Table 3).

The consistency between all models except for the General model indicates that the major reason for discrepancy is the prior on the receptor concentration. In all but the General model, the titrand concentration freely changes (subject to the constraint [*R*]_0_ > 0) from stated concentration without penalty. In the General model, the prior penalizes deviations from the stated value of [*R*]_0_. Estimates of the concentration affect Δ*H* but not Δ*G* because concentrations are highly correlated with Δ*H* but not with Δ*G*.

It is also evident that the titrand concentration is the determining factor for the shift in median Δ*H* because the titrant concentration is lognormal in all the Bayesian priors. Modifying the standard deviation in the lognormal distribution affects credible intervals but does not change the median. By elimination, the factor that leads to the shift in the median is usage of a lognormal instead of uniform prior for the titrand concentration.

**Figure 3.**
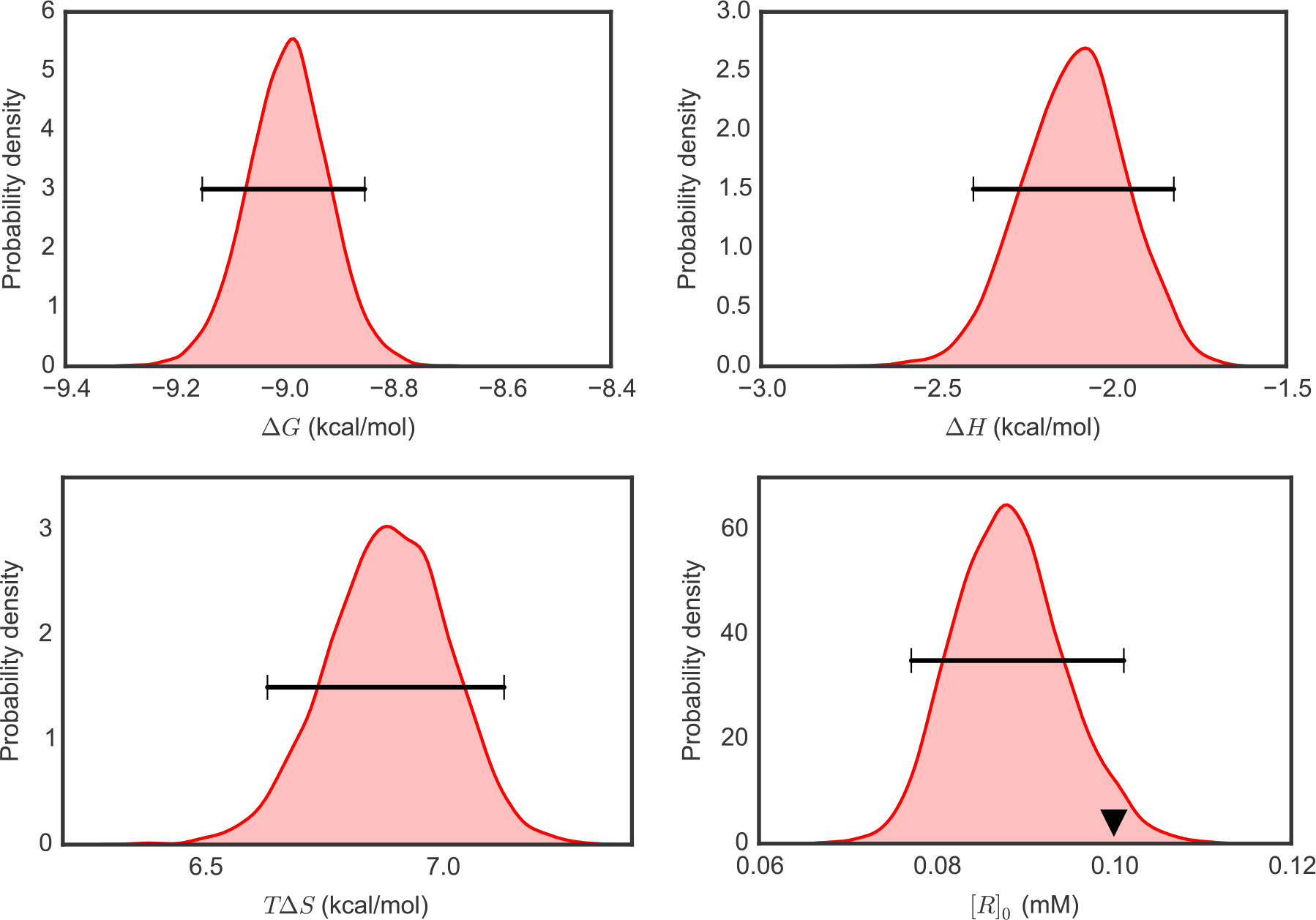
Representative 1D marginal distributions of thermodynamic parameters from Bayesian ITC analysis. 1D marginal probability densities for thermodynamic parameters of interest were estimated based on 5000 MCMC samples generated from the Bayesian posterior (General model) for one ITC experiment measuring Mg(II):EDTA binding. Horizontal bars show 95% Bayesian credible intervals. The triangle in density plot of [*R*]*_0_* indicates the stated value.

**Figure 4.**
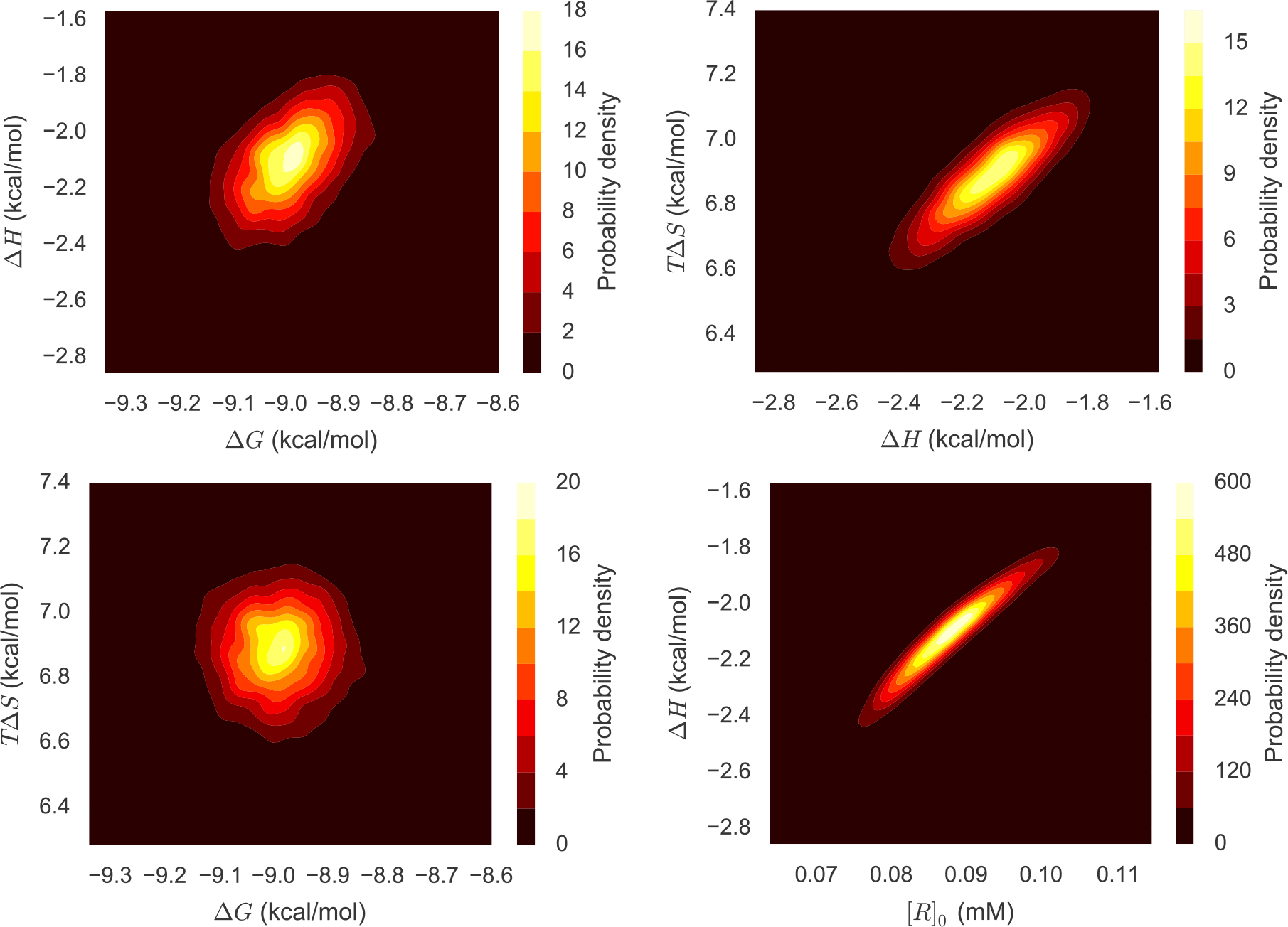
Representative 2D marginal distributions of pairs of thermodynamic properties from Bayesian ITC analysis. 2D joint marginal probability densities were estimated based on 5000 MCMC samples generated from the Bayesian posterior (General model) for one ITC experiment measuring Mg(II):EDTA binding.

**Figure 5.**
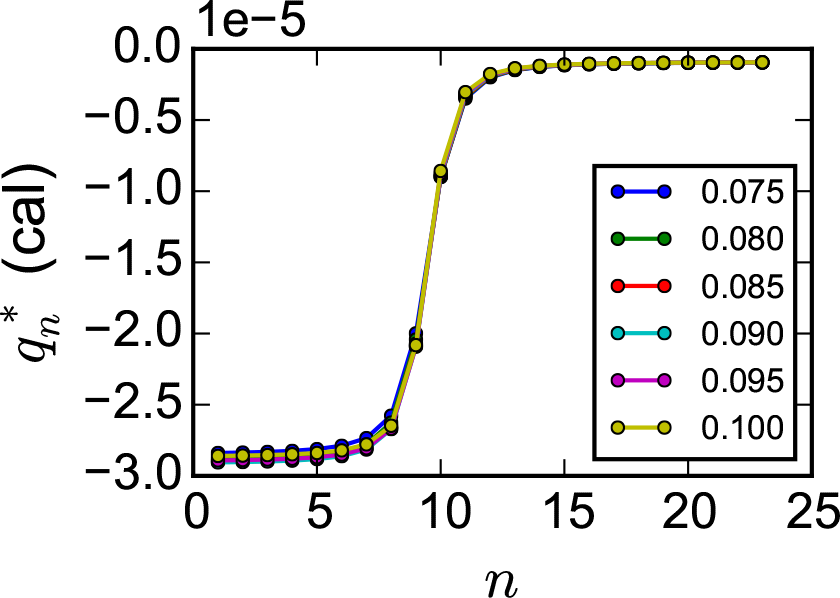
Integrated heat for different values of titrand concentration [*R*]_0_ for Mg(II):EDTA binding. Corresponding values of [*L*]_*s*_ and Δ*H* were based on a simple linear regression of [*L*]_*s*_ and of Δ*H* versus [*R*]_0_. The other parameters (Δ*G*, Δ*H*_0_) took the last value from the MCMC time series (General model).

**Table 3.** Median estimates of Δ*H* (kcal/mol). For nonlinear least squares, the value is the median of the different point estimates across different measurements. For Bayesian analysis, it is the median of the median sample from each Bayesian posterior. The numbers in parentheses are standard deviations estimated by bootstrapping: resampling the datasets (for nonlinear least squares) or the MCMC samples (for Bayesian analysis) with replacement 1000 times.

| Dataset | Nonlinear least squares | Comparison | Flat $[R]_0$ | General |
| --- | --- | --- | --- | --- |
| Mg(II):EDTA | -2.25 (0.06) | -2.24 (0.05) |  | -2.19 (0.04) |
| ligand 1:thermolysin | -5.84 (0.27) | -5.89 (0.22) | -5.92 (0.21) | -4.41 (0.31) |
| ligand 2:thermolysin | -5.08 (0.07) | -4.92 (0.06) | -4.96 (0.07) | -4.23 (0.09) |
| ligand 3:thermolysin | -5.28 (0.08) | -5.10 (0.09) | -5.14 (0.1) | -4.46 (0.29) |
| CBS:CAII | -10.36 (0.6) | -10.22 (0.53) |  | -10.05 (0.76) |

### BCIs are superior to NlRCIs

In addition to the median enthalpy, the width and and consistency of intervals is also dependent on the concentration model (See Table 4, Figure 6, and Figures S5 to S16 in the Supplementary Material). For Δ*H* and [*R*]_0_ in particular, NlRCIs and BCIs based on the Comparison model are narrower and correspondingly less consistent with one another than BCIs based on other concentration models. BCIs based on the General model are substantially broader and those based on the flat [*R*]_0_ model are broader still. However, all the BCIs and NlRCIs for Δ*G* are of comparable magnitude.

**Table 4.** Figure numbers for confidence interval plots in this manuscript.

| System | General | Flat $[R]_0$ | Comparison |
| --- | --- | --- | --- |
| Mg(II):EDTA | 6 |  | S5 |
| ligand 1:thermolysin | S6 | S7 | S8 |
| ligand 2:thermolysin | S9 | S10 | S11 |
| ligand 3:thermolysin | S12 | S13 | S14 |
| CBS:CAII | S15 |  | S16 |

In contrast with the dependence of the shift in the median enthalpy on the titrand concentration model, the change in interval size is primarily driven by the titrant concentration model. The Comparison and Flat [*R*]_0_ model have the same uniform prior for the titrand concentration. However, the size of the Δ*H* and [*R*]_0_ intervals for the Flat [*R*]_0_ model is much larger because the standard deviation in the lognormal model for [*L*]_*s*_ is larger.

On a note related to the width and consistency between confidence intervals, nearly every pair of 95% BCIs for Δ*G* and Δ*H* from the General and Flat [*R*]_0_ model have at least some overlap with one another. (The 95% BCIs for [*R*]_0_ do not overlap when the stated concentrations differ, as in the Mg(II):EDTA and CBS:CAII datasets.) As with other statistics, BCIs based on the Comparison model are very similar to NlRCIs (Figures S5, S8, S11, S14, and S16 in the Supplementary Material).

One complication with assessing confidence interval estimates is that we do not know the “true” value. Because we do not know the “true” value, we used the median value from repeated experiments as an approximation. The mean value is also a suitable choice, but the median is less sensitive to outliers.

Most of the 95% BCIs for Δ*G*, Δ*H*, and [*R*]_0_ from the General and Flat [*R*]_0_ models contain the median. One exception is for the CBS:CAII dataset, in which BCIs for Δ*G* capture the median less consistently. In contrast, while most 95% NlRCIs for Δ*G* contain the median (except in the CBS:CAII dataset), the 95% NlRCIs for Δ*H* and [*R*]_0_ generally do not. BCIs from the Comparison model behave similarly to NlRCIs (Figures S5, S8, S11, S14, and S16 in the Supplementary Material). The size of these intervals appear to be significantly underestimated in all of our systems.

**Figure 6.**
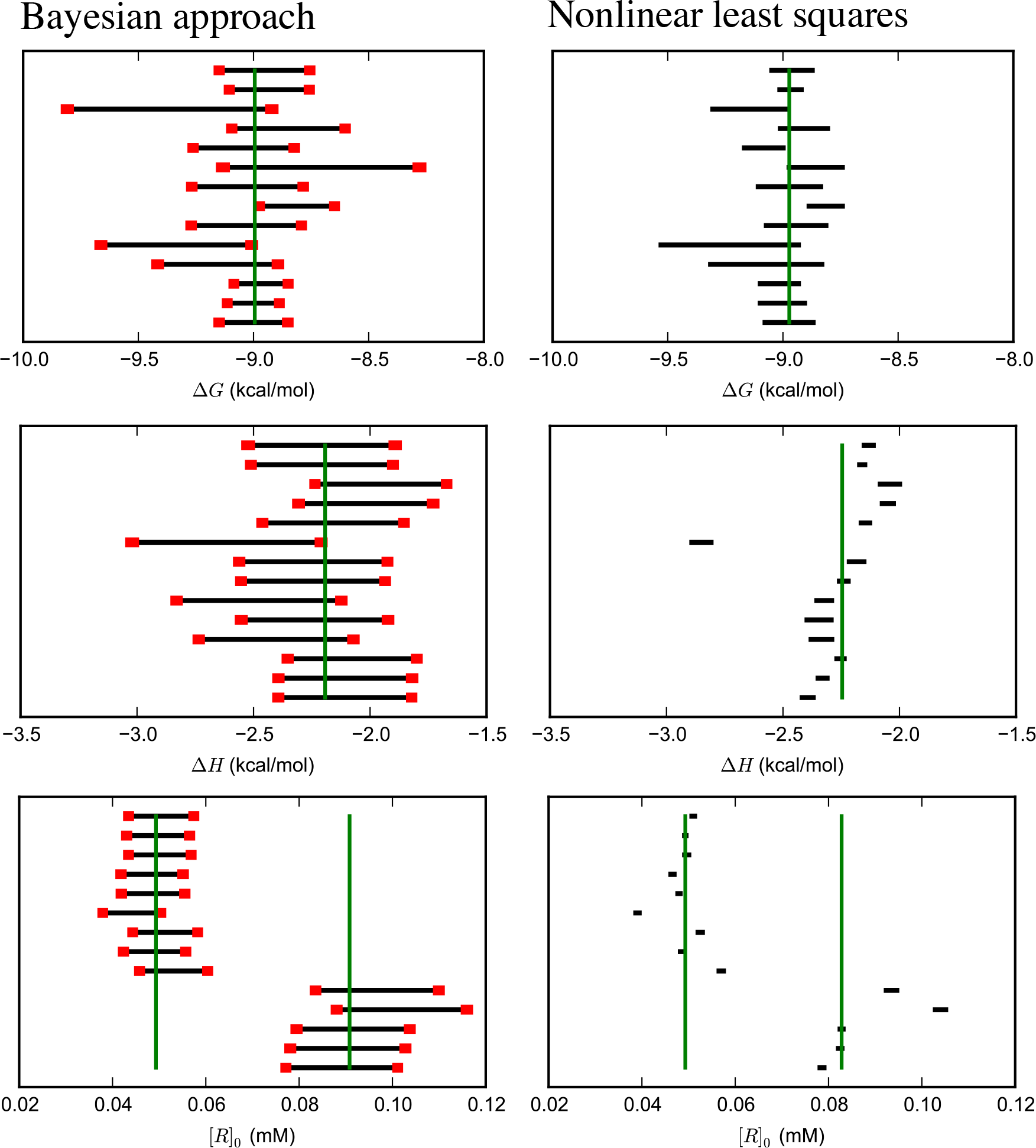
Uncertainty estimates from Bayesian and nonlinear least squares analyses of Mg(II):EDTA ITC replicates. 95% credible intervals estimated from the Bayesian posterior based on the General model (left) and confidence intervals from nonlinear least squares (right) for parameters specifying magnesium binding to EDTA. The vertical green lines are the median. There are two median estimates for *R* because the experiments were done at two different concentrations. Red bars denote the standard deviations of the lower and upper bounds, estimated by bootstrapping, and are a total of two standard deviations wide.

A better way to visualize the performance of BCIs and NlRCIs as confidence intervals is to compare the fraction of intervals that contain the median with the stated confidence level. If the confidence levels truly specify the probability of containing the true value, then data points should lie along the diagonal solid line of Figure 7 and Figures S17 to S20 in the Supplementary Material. Points below the diagonal indicate that stated confidence intervals are too small; points above the diagonal indicate that they are too large.

By this metric, BCIs based on the General model perform nearly ideally for Mg(II):EDTA and less reliably for the other datasets. In the cases of Mg(II):EDTA and ligand **2**:thermolysin binding, the observed fraction of BCIs (General model) for Δ*G* and Δ*H* that contain the median is very close to the ideal line (Figure 7 and Figure S18 in the Supplementary Material). For the other datasets, BCIs based on the General model are less consistent with observed rates. In the cases of ligand **1**:thermolysin and ligand **3**:thermolysin, the median-containing frequency of Δ*G* BCIs is also very close the ideal line whereas that Δ*H* BCIs deviates from ideality, especially for larger confidence intervals (Figures S17 and S19 in Supplementary Material). In the CBS:CAII dataset, however, BCIs for Δ*H* are more consistent with observed rates than for Δ*G*.

Intervals from other models had variable performance. NlRCIs of Δ*G* have similar performance to BCIs but the observed rate at which NlRCIs for Δ*H* contain the median is significantly less than ideal. This deviation from ideality is consistent with the poorly overlapping 95 % confidence intervals for Δ*H*. BCIs from the Comparison model behave similarly to NlRCIs. In contrast with intervals from other models, BCIs based on the flat [*R*]_0_ model generally overestimate the width of intervals for the thermolysin model. The overestimation of intervals suggests that the uniform prior employed in this analysis is too uninformative.

Overall, our Bayesian method (with the General model) led to reasonable BCIs for multiple measurements performed by a single individual within a single laboratory. The performance of BCIs in accounting for laboratory-to-laboratory variability in the CBS:CAII datasets digitized from the ABRF-MIRG’02 paper [21] was weaker. In this dataset, there must be one or more significant sources of error that the present approach fails to account for.

The strong correlation between concentrations and Δ*H* explains the dramatic improvement of the credible intervals of Δ*H* (e.g. Figure 7) when the uncertainty in [*L*]_*s*_ is included in the Bayesian analysis. In the same vein, the weak correlation between concentrations and Δ*G* explains why NlRCIs for Δ*G* are reasonable (Figure 7) even if the titrant concentration was treated as exactly known in the fit. Trends in the accuracy of confidence intervals are consistent with previous analyses based on error propagation [1, 5, 30, 34], which showed that titrant concentration errors propagate to small relative errors in Δ*G* but large relative errors in Δ*H*. If the error in titrant concentration is correctly propagated, it may be possible to make NlRCIs more accurate [1], but testing this is beyond the scope of the present work. In subsequent analysis, we will only consider the General model.

**Figure 7.**
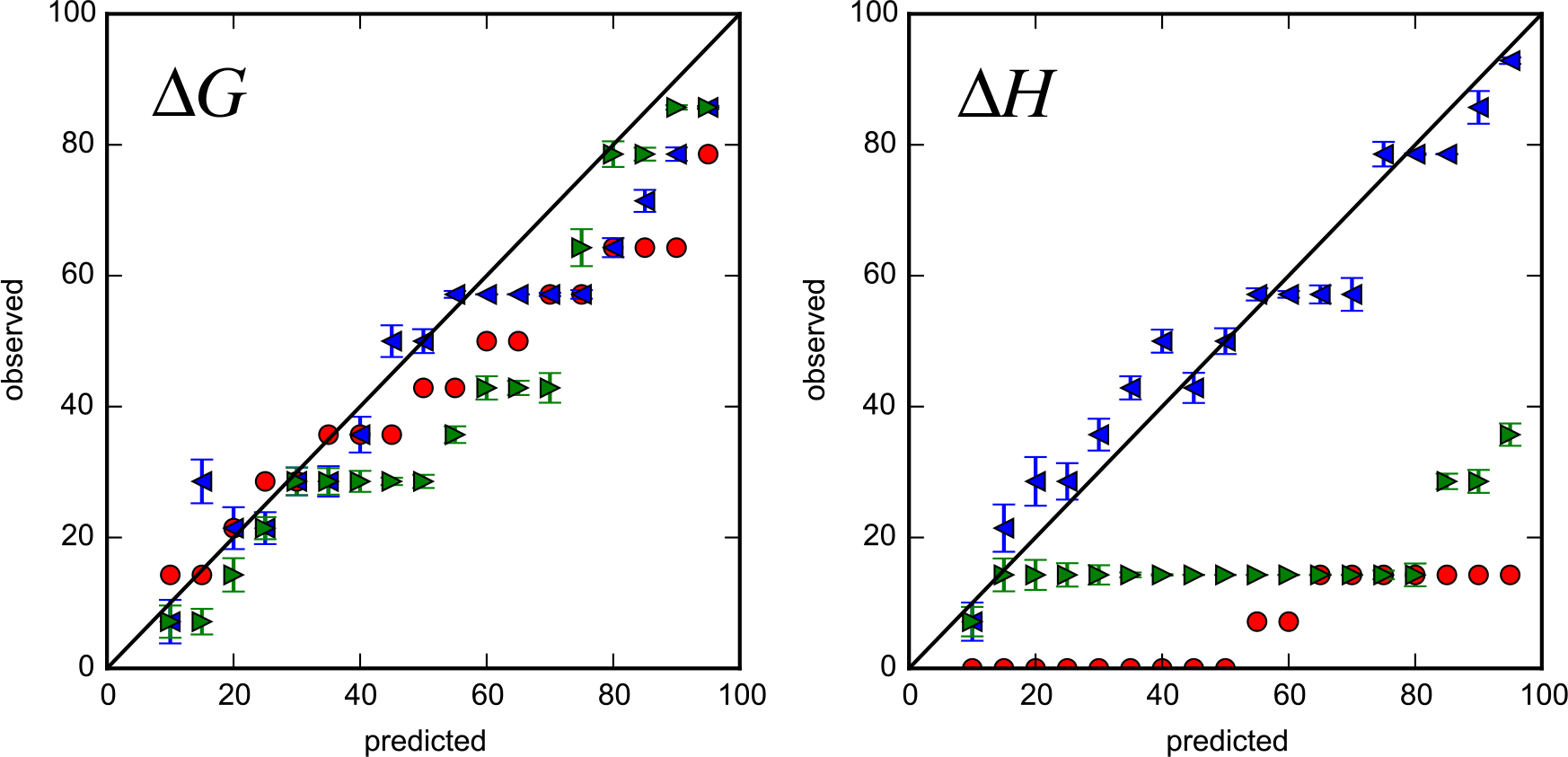
Uncertainty validation for Bayesian and nonlinear least squares analyses of Mg(II):EDTA data. For the Mg2:EDTA binding experiments, the predicted versus observed rate (%) in which intervals contain the median value for binding parameters is shown. Intervals were BCIs based on the General model (blue leftward triangles), Comparison model (green rightward triangles), or nonlinear least squares confidence intervals (red circles). Error bars are standard deviations based on bootstrapping.

### Binding parameter distributions are more consistent with Bayesian analysis than nonlinear regression

In most datasets, the estimated Kullback-Leibler divergence between pairs of Bayesian posteriors is smaller than those estimated for nonlinear regression (Figure 8 and Figures S21 to S24 in Supplementary Material.) For the thermolysin datasets where the fiat [*R*]_0_ model was tested, the Kullback-Leibler divergence for the flat [*R*]_0_ model was even smaller than for the general model. Therefore, marginals of the Bayesian posteriors are more consistent with one another than the Gaussian distributions from nonlinear regression. This finding agrees with above analyses that the Bayesian posterior captures the variance among experiments better than nonlinear least squares. The one exception is with the CBS:CAII dataset, in which the Kullback-Leibler divergence matrix based on the Bayesian method is comparable to the one from nonlinear regression (Figure S21 in the Supplementary Material).

**Figure 8.**
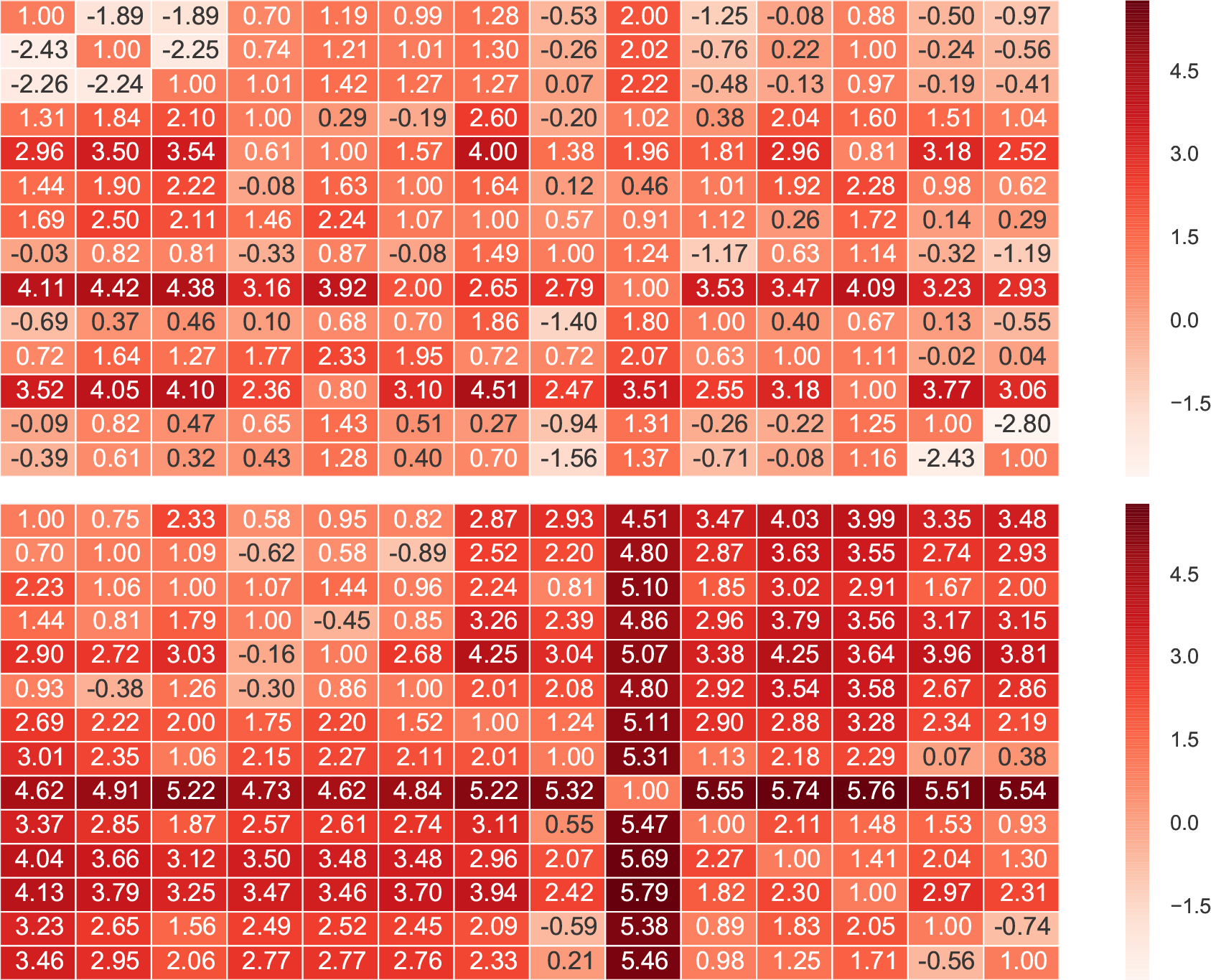
The natural logarithm of Kullback-Leibler divergence between Bayesian approach (General model) and nonlinear least-squares. The natural logarithm of the KL-divergence between posterior marginal distributions (top) and between Gaussian distributions of nonlinear least squares errors (bottom) is shown. Each column and row corresponds to one of the 14 datasets of Mg(II):EDTA binding. The diagonal elements should be ln0 = −∞ but were set to 1 for visualization.

## Conclusions

In this study we have applied Bayesian statistics to analyze ITC data for the first time. We were able to account for various sources of error including, most importantly, uncertainties in the titrand and titrant concentrations. Due to the inclusion of concentration uncertainties, BCIs more accurately capture the 15 of 21variance between independent experiments than NlRCIs. In some datasets, the concentration error model led to differences in binding enthalpy estimates. Correlation between different parameters computed from the Bayesian posterior helps rationalize the effects of concentration uncertainty on the accuracy of Δ*G* and Δ*H*.

## Acknowledgments

We thank Gerhard Klebe for facilitating the sharing of ITC data collected by his former student Stefan Krimmer. We also thank Joel Tellinghuisen for helpful discussions and comments on the manuscript. JDC thanks Sarah Boyce for training on ITC instruments. Finally, we thank our summer interns at Illinois Tech: Mateus Pires Schneider (funded by the Capes Foundation within the Brazilian Ministry of Education) for performing NITPIC integrations on the thermolysin systems; and Erica Cusnariov for assistance with KL divergence figures. ASR and JDC acknowledge support from the Sloan Kettering Institute. JDC acknowledges support from NIH grant P30 CA008748 and NIH grant R01 GM121505. THN and DDLM acknowledge support from NIH grant R15 GM114781.

## Appendix Simple two-component (1:1) association binding model

In a simple two-component (1:1) complexation reaction, we have reversible association between a ligand *L* and a receptor *R*

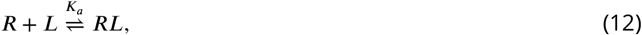

where the *association constant K_a_* or the binding free energy Δ*G* is related to concentrations [*X*] of the microscopic species at equilibrium by,

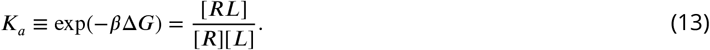

With each injection, three effects will contribute to the true quantity of heat 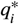 liberated due to injection *i*: (1) the association of *R* with *L*, (2) the dilution of ligand and buffer into the protein solution (as most solutions are nonideal), and (3) the mechanical heat produced by the injection and stirring. We subsume the latter two components into a single term Δ*H*_0_, and write

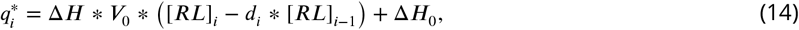

where Δ*H* is the enthalpy change associated with binding, [*RL*]_*i*_ is the complex equilibrium concentration after injection *i*, *V*_0_ is the cell volume, and *d*_*i*_ is the dilution factor after an injection with volume *v*_*i*_, defined as

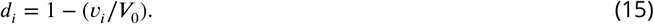

In what follows, we will express the complex equilibrium concentration after injection *i*, [*RL*]_*i*_, in terms of *K*_*a*_, the cell volume *V*_0_, the initial concentration of the receptor [*R*]_0_, and syringe concentration of the ligand [*L*]_*s*_.

The total quantity (number of moles) of receptor *R*_*i*_ and ligand *L*_*i*_ in the cell after injection *i* is given by

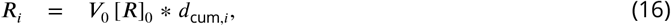

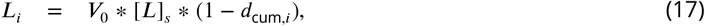

where *d*_cum,*i*_ is the cumulative dilution factor given by

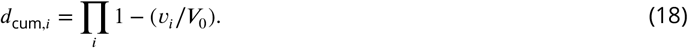

This model accounts for perfusion of receptor and ligand from the cell at a constant cell volume while assuming the mixing after injection is instantaneous as in [32].

Conservation of mass gives us the constraints

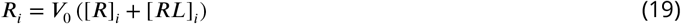

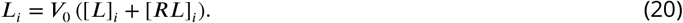

Combining Eqs. 13 and 20 gives

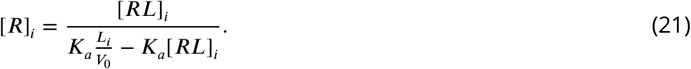

Substituting Eq. 21 into Eq. 19 yields a quadratic equation in the complex equilibrium concentration [*RL*]_*i*_:

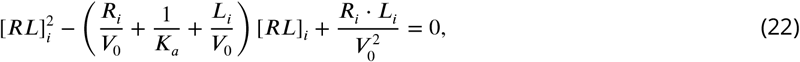

where the only solution that satisfies 0 ≤ [*RL*]_*i*_ ≤ min{[*R*], [*L*]_*i*_} is

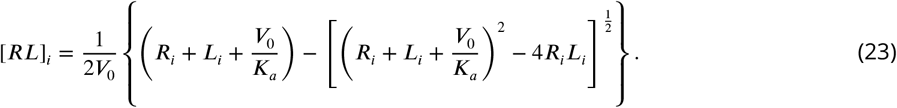

